# Simple statistical identification and removal of contaminant sequences in marker-gene and metagenomics data

**DOI:** 10.1101/221499

**Authors:** Nicole M. Davis, Diana M. Proctor, Susan P. Holmes, David A. Relman, Benjamin J. Callahan

**Affiliations:** Department of Microbiology and Immunology, Stanford University School of Medicine, Stanford, CA 94305, USA; Department of Medicine, Stanford University School of Medicine, Stanford, CA 94305, USA; Department of Orofacial Sciences, University of California, San Francisco School of Dentistry, San Francisco, CA 94143, USA; Department of Statistics, Stanford University, Stanford, CA 94305, USA; Infectious Diseases Section, Veterans Affairs Palo Alto Health Care System, Palo Alto, CA 94304, USA; Department of Population Health and Pathobiology, College of Veterinary Medicine, North Carolina State University, Raleigh, NC 27607, USA; Bioinformatics Research Center, North Carolina State University, Raleigh, NC 27695, USA

**Author notes:** Equal contributions (Nicole M. Davis, Diana M. Proctor). Equal contributions (David A. Relman, Benjamin J. Callahan). **Correspondence:** 456 Research Building College of Veterinary Medicine 1060 William Moore Drive Raleigh, NC 27607.

**Keywords:** microbiome, metagenomics, marker-gene, 16S rRNA gene, DNA contamination

## Abstract

**Background:** The accuracy of microbial community surveys based on marker-gene and metagenomic sequencing (MGS) suffers from the presence of contaminants — DNA sequences not truly present in the sample. Contaminants come from various sources, including reagents. Appropriate laboratory practices can reduce contamination, but do not eliminate it. Here we introduce `decontam` (https://github.com/benjjneb/decontam), an open-source R package that implements a statistical classification procedure that identifies contaminants in MGS data based on two widely reproduced patterns: contaminants appear at higher frequencies in low-concentration samples, and are often found in negative controls.

**Results:** `decontam` classified amplicon sequence variants (ASVs) in a human oral dataset consistently with prior microscopic observations of the microbial taxa inhabiting that environment and previous reports of contaminant taxa. In metagenomics and marker-gene measurements of a dilution series, `decontam` substantially reduced technical variation arising from different sequencing protocols. The application of `decontam` to two recently published datasets corroborated and extended their conclusions that little evidence existed for an indigenous placenta microbiome, and that some low-frequency taxa seemingly associated with preterm birth were contaminants.

**Conclusions:** `decontam` improves the quality of metagenomic and marker-gene sequencing by identifying and removing contaminant DNA sequences. `decontam` integrates easily with existing MGS workflows, and allows researchers to generate more accurate profiles of microbial communities at little to no additional cost.

## Background

High-throughput sequencing of DNA from environmental samples is a powerful tool for investigating microbial and non-microbial communities. Community composition can be characterized by sequencing taxonomically informative marker genes, such as the 16S rRNA gene in bacteria [1–4]. Shotgun metagenomics, in which all DNA recovered from a sample is sequenced, can also characterize functional potential [5–7]. However, the accuracy of marker-gene and metagenomic sequencing (MGS) is limited in practice by several processes that introduce contaminants — DNA sequences not truly present in the sampled community.

Failure to account for DNA contamination can lead to inaccurate data interpretation. Contamination falsely inflates within-sample diversity [8, 9], obscures differences between samples [8, 10], and interferes with comparisons across studies [10, 11]. Contamination disproportionately affects samples from low-biomass environments with less endogenous sample DNA [10, 12–16], and can lead to controversial claims about the presence of bacteria in low microbial biomass environments like blood and body tissues [12, 13, 15–17]. In high-biomass environments, contaminants can comprise a significant fraction of low-frequency sequences in the data [18], limiting reliable resolution of low-frequency variants and contributing to false-positive associations in exploratory analyses [19].

Attempts to control DNA contamination before and after sequencing have had mixed success. One common practice is to process reagent-only [9,14, 20] or blank sampling instrument [21] negative control samples alongside biological samples at the DNA extraction and PCR steps. Contamination is often assumed to be absent if control samples do not yield a band on an agarose gel [22, 23]. However, band-less samples can generate non-negligible numbers of sequencing reads [14, 15], suggesting that gel-based quality control is insufficient.

There are two major types of contaminants in MGS experiments that arise from different sources. External contamination is contributed from outside the samples being measured, with potential sources that include research subjects’ or investigators’ bodies [24, 25], laboratory surfaces and air [10,21,26], and, perhaps most importantly, sample collection instruments and laboratory reagents [9,12,14]. Internal or cross-contamination arises when samples mix with each other during sample processing [9] or sequencing [33]. Contamination can be reduced through laboratory techniques such as UV irradiation, “ultrapurification” and/or enzymatic treatment of reagents, and the separation of pre- and post-PCR areas [9,12,27,28]. However, even optimal lab practices do not completely eliminate DNA contamination [11, 14].

*In silico* contaminant removal can complement existing laboratory approaches, but distinguishing contaminating microbial DNA from true microbial sequences can be difficult and is not often performed [14, 16]. Perhaps the most common *in silico* decontamination method in practice is the removal of sequences below an *ad hoc* abundance threshold [21,29–31]. However, abundance thresholds remove rare features truly present in the sample, and do not remove abundant contaminants that are the most likely to interfere with subsequent analysis. Another approach is the removal of sequences that appear in negative controls [e.g. 10, 32, 20]. However, cross-contamination between samples often causes abundant true sequences to be detected in negative controls [9, 19, 33]. Finally, “blacklist” methods exclude sequences or taxa previously identified as contaminants, but do not identify study-specific contaminants and often remove true sequences.

Despite the widespread problem of contamination, few software tools exist that directly address MGS contaminants. SourceTracker uses Bayesian mixtures to identify the proportion of a sample consistent with origin from external contaminating sources of known composition, but does not identify specific contaminants [26]. The visualizations and summary statistics provided by the An’vio software package can be used to identify contaminant metagenome-assembled-genomes (MAGs) [34], but this relies on user expertise to identify contaminant-specific patterns and does not apply to marker-gene data. Recently, a new method for identifying cross-contaminants arising from index switching during sequencing has been developed for dual-indexed MGS libraries, but this method does not apply to other types of contamination [33].

Here, we introduce and validate decontam, a simple-to-use open-source R package that identifies and removes external contaminants in MGS data. `decontam` implements two simple *de novo* classification methods based on widely reproduced signatures of external contamination: 1) Sequences from contaminating taxa are likely to have *frequencies* that inversely correlate with sample DNA concentration [8,14,16,30], and 2) sequences from contaminating taxa are likely to have higher *prevalence* in control samples than in true samples [10,31,32]. Frequency-based contaminant identification relies on auxiliary DNA quantitation data that is in most cases intrinsic to MGS sample preparation. Prevalence-based contaminant identification relies on sequenced negative controls [11, 14]. `decontam` is not intended to detect cross-contamination, which presents with qualitatively different statistical patterns in MGS data.

We validated `decontam` on marker-gene and metagenomics datasets generated by our laboratory and others. In an oral 16S rRNA gene dataset, `decontam` classifications were consistent with curated reference databases of common contaminating microbial genera and known oral microbes. In data generated by Salter *et al.*, `decontam` selectively removed contaminants, thereby reducing technical variation due to sequencing center or DNA extraction kit in marker-gene and shotgun metagenomics data, respectively. The application of `decontam` to 16S rRNA gene sequencing data generated from placenta biopsies corroborated the conclusion that the data did not support the existence of a placenta microbiome [15]. `decontam` improved a recent exploratory analysis of associations between preterm birth and the vaginal microbiota by identifying run-specific contaminants [19]. Our results suggest that `decontam` distinguishes contaminants from non-contaminants across diverse studies, and that removal of these contaminants improves the accuracy of biological inferences in studies that use MGS methods to investigate microbial communities.

## Description of the Method

### Frequency-based contaminant identification

Let total sample DNA (T) be a mixture of two components (T = C + S): contaminating DNA (C) present in uniform concentration across samples, and true sample DNA (S) present in varying concentration across samples. Let the frequency of a sequence, or set of sequences, be its abundance divided by the total abundance of all sequences in the sample (alternative terms used equivalently include proportion and relative abundance). In the limit S >> C, the frequency of contaminating DNA (f_C_) is inversely proportional to total DNA T (Fig. 1), while the frequency of sample DNA (f_S_) is independent of T:

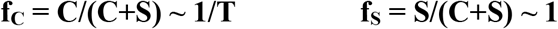

**Figure 1.**
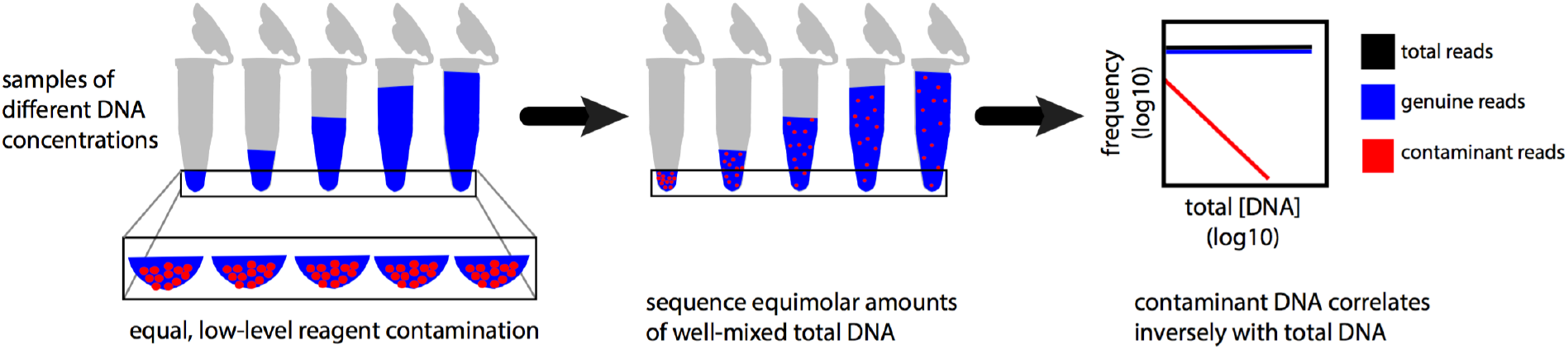
Mixture model of contaminants and non-contaminants in MGS experiments. Contaminant DNA is expected to be present in approximately equal and low concentrations across samples, while sample DNA concentrations can vary widely. As a result, the expected frequency of contaminant DNA varies inversely with total sample DNA concentration (red), while the expected frequency of non-contaminant DNA does not (blue).

For each sequence feature two models are compared: a contaminant model, in which expected frequency varies inversely with total DNA concentration, and a non-contaminant model, in which expected frequency is independent of total DNA concentration. More precisely, two linear models are fit to the log-transformed frequencies as a function of the log-transformed total DNA, a contaminant model with slope −1 and a non-contaminant model with slope 0. Samples in which the sequence feature is absent are omitted. The ratio R between the sums-of-squared-residuals of the contaminant and non-contaminant models is computed, and then the score statistic P is defined as the tail probability at value R of an F distribution with degrees of freedom equal to the number of samples in which the feature was present. The score statistic P ranges from 0 to 1. Small scores indicate the contaminant model is a better fit, and high scores indicate that the non-contaminant model is a better fit.

Although inspired by the general linear F test, the frequency-based score statistic is not a p-value, i.e. it is not associated with any guarantees on the Type 1 error rate. The frequency-based score statistic is best thought of as a transformation that takes as input two values — the ratio of the sum-of-squared-residuals, and the number of observations — and outputs a score that is a better classification statistic than the ratio of the sum-of-squared-residuals alone. This transformation differentiates between otherwise identical ratios of sum-of-squared-residuals that are supported by different numbers of observations, and appropriately outputs scores closer to the extremes (0/1) when non-unitary ratios are supported by more observations.

Frequency-based contaminant identification is not recommended for extremely low-biomass samples (C ~ S or C > S) because the simple approximations we are making for the dependence of contaminant frequency on total DNA concentration break down when contaminants comprise a large fraction of sequencing reads.

### Prevalence-based contaminant identification

Once again, let total sample DNA (T) be a mixture of contaminating DNA (C) and true sample DNA (S), i.e. T = C + S. The results of MGS sequencing can be thought of as an incomplete sampling of T. Thus, in negative controls (S ~ 0) the likelihood of detecting any given contaminant sequence feature will be higher than in true samples (S > 0). That is, the prevalence of contaminants will be higher in negative controls than in true samples due to the absence of competing DNA in the sequencing process.

For each sequence feature, a chi-square statistic on the 2×2 presence-absence table in true samples and negative controls is computed, and a score statistic P is defined as the tail probability of the chi-square distribution at that value. The p-value from Fisher’s exact test is used as the score statistic instead if there are too few samples for the chi-square approximation. The score statistic ranges from 0 to 1. Small scores indicate the contaminant model of higher prevalence in negative control samples is a better fit.

Although the prevalence-based score statistic is set equal to a p-value from the chi-square or Fisher’s exact tests, it is used by `decontam` only as a score that effectively distinguishes between the contaminant and non-contaminant mixture components. This treatment is also recommended by the potential for cross-contamination to violate distributional assumptions related to independence between samples.

We recommend use of the prevalence method in very low biomass environments where a majority of MGS sequences might derive from contaminants rather than true inhabitants of the sampled environment (i.e. C ~ S or C > S). Even in the low-biomass regime, it is still expected that non-contaminants will appear in a larger fraction of true samples than in negative control samples.

### Sequencing Batches and Composite Identification

Separately processed samples may have different contaminants [12,14,28,36]. `decontam` allows the user to specify processing batches, in which case score statistics are generated from each batch independently and then combined in a user-selectable fashion for classification (for example, by taking the minimum score across batches). `decontam` also provides simple methods to combine scores from the frequency and prevalence methods into a composite score statistic. For example, the combined method uses Fisher’s method to combine the frequency and prevalence score statistics (interpreted as tail probabilities) into a composite score statistic that is then used for classification.

### Classification

A sequence feature is classified as contaminant or non-contaminant by comparing its associated score statistic P to a user-defined threshold P*, where P can be the frequency, prevalence, or composite score. If P < P*, the sequence feature is classified as a contaminant. The default classification threshold is P*=0.1, but we highly recommend that users inspect the distribution of scores in their data and consider adjusting P* based on specific dataset characteristics (see also Discussion). The threshold P*=0.5 has a particularly simple interpretation: In the frequency approach, sequence features would be classified as contaminants if the contaminant model is a better fit than the non-contaminant model, and in the prevalence approach sequence features would be classified as contaminants if present in a higher fraction of negative controls than true samples.

### The decontam R package

The contaminant classification methods introduced here are implemented in the open-source `decontam` R package available from GitHub (https://github.com/benjjneb/decontam) and the Bioconductor repository [35]. The primary function, *isContaminant*, implements frequency- and prevalence-based contaminant identification that can be applied to variety of sequence features including amplicon sequence variants (ASVs), operational taxonomic units (OTUs), taxonomic groups (e.g. genera), orthologous genes, metagenome-assembled-genomes (MAGs), and any other feature with quantitative per-sample abundance that is derived from marker-gene or metagenomics sequencing data (see also Discussion).

The primary input to *isContaminant* is a feature table of the abundances or frequencies of sequence features in each sample (e.g. an OTU table). In addition, *isContaminant* requires one of two types of auxiliary data for frequency- and prevalence-based contaminant identification, respectively: 1) Quantitative DNA concentrations for each sample, often obtained during amplicon or shotgun sequencing library preparation in the form of a standardized fluorescence intensity (e.g. PicoGreen), and/or 2) sequenced negative control samples, preferably DNA extraction controls to which no sample DNA was added. Contaminants identified by `decontam` can be removed from the feature table with basic R functions described in `decontam` vignettes.

The *isNotContaminant* function supports the alternative use case of identifying noncontaminant sequence features in very low biomass samples (C > S). *isNotContaminant*implements the prevalence method, but with the standard prevalence score P replaced with 1-P, so low scores are now those associated with non-contaminants. *isNotContaminant* does not implement the frequency method for reasons described above, and classifies very low prevalence samples conservatively, i.e. as contaminants, as is appropriate for the low-biomass regime.

## Results

### decontam discriminates likely contaminants from likely inhabitants of the human oral cavity

As part of an ongoing study of the human oral microbiome [37], we processed 33 reagent-only or blank-swab DNA extraction negative control samples alongside and in the same manner as 712 oral mucosa samples. We inspected the frequencies of amplicon sequence variants (ASVs) as a function of DNA concentration (range: undetectable-39 ng/uL) measured by fluorescent intensity after PCR and prior to sequencing. Two clear patterns emerged (Fig. 2a): ASV frequencies that were independent of DNA concentration, and ASV frequencies that were inversely proportional to DNA concentration [8,14,16,30], consistent with total DNA consisting of a mixture of contaminant and non-contaminant components. Taxonomic assignments for ASVs with inverse frequency patterns were consistent with contamination. For example, Seq3 was a fungal mitochondrial DNA sequence, while Seq53 and Seq152 were assigned to the commonly contaminating genera *Methylobacterium* and *Phyllobacterium* [8,12,14,15]. Taxonomic assignments of ASVs with frequencies independent of sample DNA concentration were consistent with membership in the oral microbiota. For example: Seq1, *Streptococcus sp.*, Seq12, *Neisseria sp.*, and Seq200, *Treponema sp*. [37–41]. The total concentration of contaminants assigned by the prevalence method was roughly constant and independent of total DNA concentration, consistent with our mixture model (Fig. S1).

**Figure 2.**
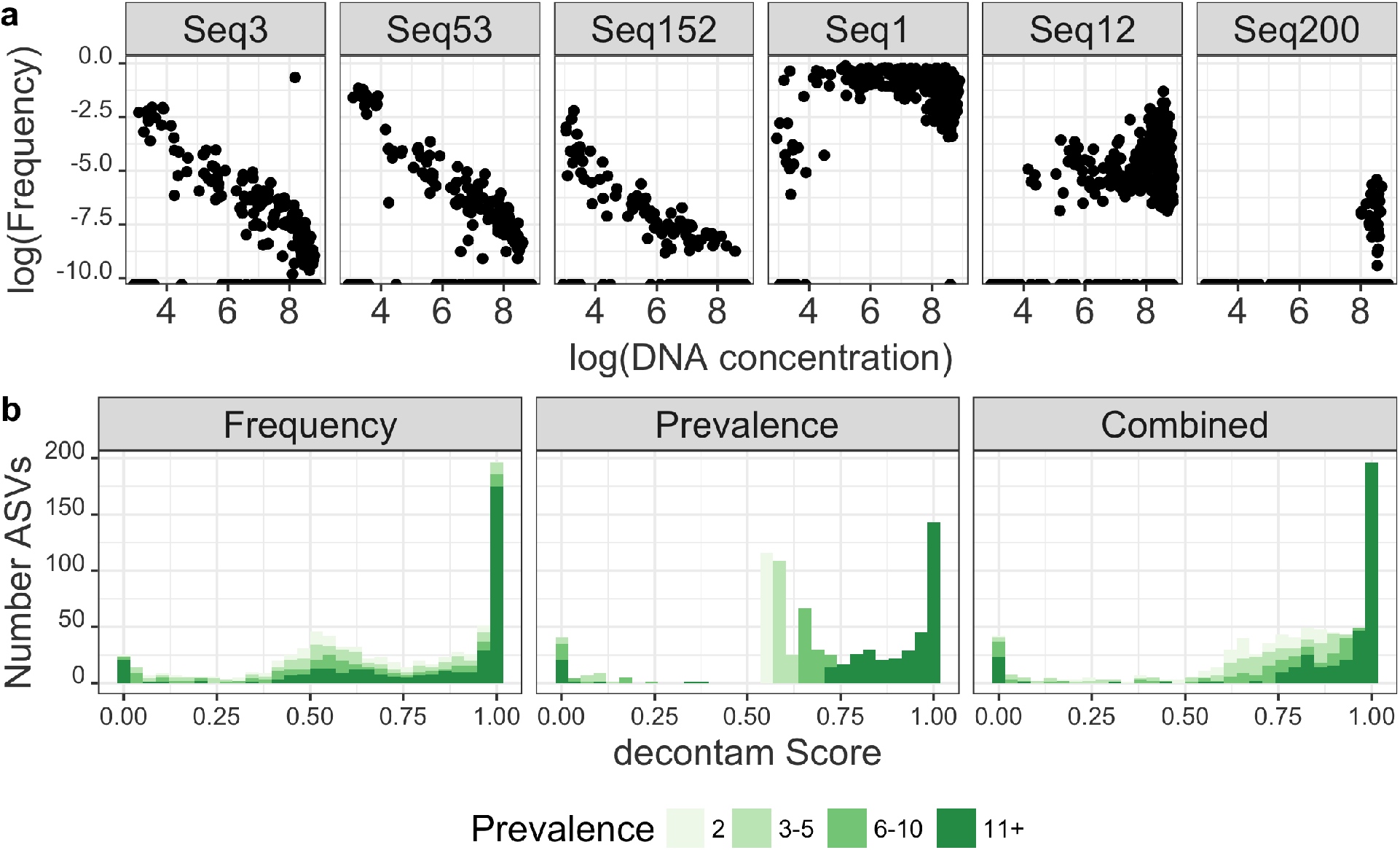
Frequency patterns and decontam score distributions of microbial sequences from an oral mucosal 16S rRNA gene dataset. **(a)** Frequency patterns of six sequences from a 16S rRNA gene study of human oral microbial communities. The frequencies of sequence variants Seq3, Seq152, and Seq53 vary inversely with sample DNA, a characteristic of contaminants. The frequencies of Seq1, Seq12, and Seq200 are independent of sample DNA concentration, a characteristic of genuine sample sequences. **(b)** Scores for each amplicon sequence variant (ASV) present in two or more samples were computed by the frequency, prevalence and combined methods as implemented in the *isContaminant* function in the `decontam` R package. The histogram of scores is shown, with color intensity depending on the number of samples (or prevalence) in which each ASV was present.

The distribution of scores assigned by `decontam` reflected the bimodal distribution expected from a mixture of contaminant and non-contaminant components (Fig. 2b), albeit with an additional mode near 0.5 consisting of low-prevalence taxa for which `decontam` has little discriminatory power. Most ASVs (Fig. 2b) and an even larger majority of total reads (Fig. S2) were assigned high scores suggesting non-contaminant origin. The distribution of scores from the combined method, which combines the frequency-based and prevalence-based scores into a composite score, had the cleanest bimodal distribution (Fig. 2b), suggesting it will provide the most robust classifications when both DNA concentration and negative control data are available.

To assess the classification accuracy of decontam, we generated two databases to serve as proxies for true oral sequences and contaminants (Methods). `decontam` assigned scores less than 0.5 to most ASVs from genera present in the contamination database, including features Seq3, Seq53, and Seq152 (Fig. 2a). In contrast, most ASVs from genera present in the oral database, including Seq1, Seq12, and Seq200 were assigned scores greater than 0.5 (Fig. 2a). ASVs belonging to genera found in both databases or neither database display a range of scores (Fig. S3), suggesting that the reference databases constructed here incompletely separate oral and contaminating genera.

To quantitatively assess the accuracy of `decontam` we examined a restricted set of genera that were clearly and unambiguously classified as contaminants or oral taxa by our reference databases (Methods). The scores assigned by the frequency and prevalence methods to all ASVs are shown in Fig. 3A, with points colored if their genus has an unambiguous reference classification. Each panel represents a different sample prevalence threshold (e.g., whether the ASV was detected in 2, 3-5, 6, or 11+ samples). As expected, the power of `decontam` to discriminate between contaminants and non-contaminants increases with the number of samples in which each ASV was present (its prevalence). At high prevalence the frequency, prevalence and reference-based classifications are nearly identical.

**Figure 3.**
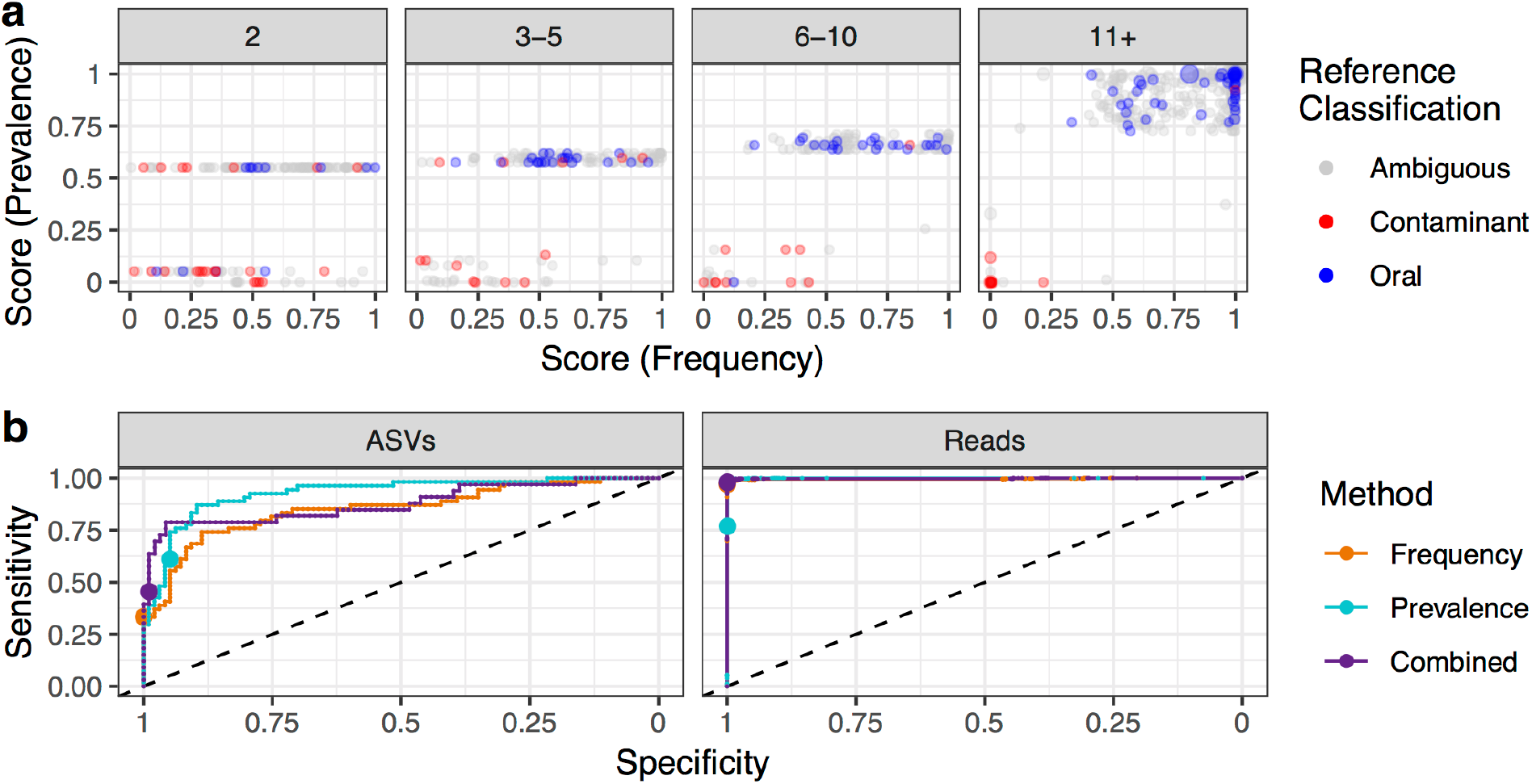
Classification accuracy of decontam on a subset of microbial sequences in an oral mucosa dataset belonging to contaminant and oral genera. **(a)** The scores assigned by the frequency and prevalence methods are plotted for each ASV present in two or more samples in the oral mucosa dataset. Points are colored if their genus can be unambiguously classified as oral or contaminant by comparison to a compiled reference database. **(b)** Receiver-operator-characteristic (ROC) curves are plotted for the frequency, prevalance and combined methods, and when evaluating sensitivity/specificity at the level of ASVs or at the level of reads (i.e. weighting by ASV abundance). Points show the default classification threshold of P*=0.1.

We developed receiver-operator-characteristic (ROC) curves of the classifier performance of the frequency, prevalence and combined methods on the subset of ASVs with unambiguous reference-based classifications, and using the reference-based classification as the ground truth. The sensitivity of all methods to detect contaminant ASVs reaches substantial levels before significant degradation in specificity occurs (Fig. 3b). At the default threshold of 0.1 (indicated by points in Fig. 3b) the frequency method has an ASV sensitivity/specificity of 0.33/ 1.00, the prevalence method 0.61/ 0.95, and the combined method 0.45/0.99. Accuracy is far higher when accounting for the abundance of each ASV and evaluating performance on a per-read basis. At the default classification threshold of 0.1, the per-read sensitivity/specificity is 0.97/1.0000 for the frequency method, 0.77/0.9994 (prevalence) and 0.98/0.9999 (combined).

These results indicate that `decontam` has high classification accuracy on the abundant and prevalent sequence features that will most impact subsequent analysis. The classification accuracy of `decontam` may be even higher than reported here given our reliance on imperfect taxonomic assignments to set a ground truth. For example, the apparent high-prevalence false-negative (the red point in the upper-right of the 11+ panel in Fig. 3a) was assigned to the genus *Peptococcus*, which is known to colonize the human mouth [42, 43]. However, *Peptococcus* was not present in our incomplete database of cultivated oral genera, so it was treated as a ground truth contaminant in our accuracy analysis.

### Application of decontam to a dilution-series test dataset

Salter *et al*. characterized a dilution series of a *Salmonella bongori* monoculture over a range of six 10-fold dilutions by 16S rRNA gene sequencing at three sequencing centers, and by shotgun metagenomics using four DNA extraction kits (one of which yielded little DNA and is excluded). Standard DNA quantitation data were not reported, so the reported 16S qPCR results (Fig. 2 from Ref. 14) were used to quantify sample DNA concentrations.

Over 50% of the contaminant (i.e. non *S. bongori)* amplicon reads and over 80% of the contaminant shotgun reads were correctly classified as contaminants by decontam’s frequency method at the default threshold P*=0.1, and sensitivity increased with higher values of P* (Fig. 4). A smaller fraction of the unique sequence features (ASVs in the 16S rRNA gene data and genera in the shotgun data) were identified as contaminants due to the small number of samples in this dataset (six samples per batch) and the lower sensitivity of `decontam` on contaminants present in few samples (e.g. Fig 3a). Identifying contaminants on a per-batch basis was more effective than pooling data across sequencing centers and DNA extraction kits, which had different contaminant spectra (Fig. 4).

**Figure 4.**
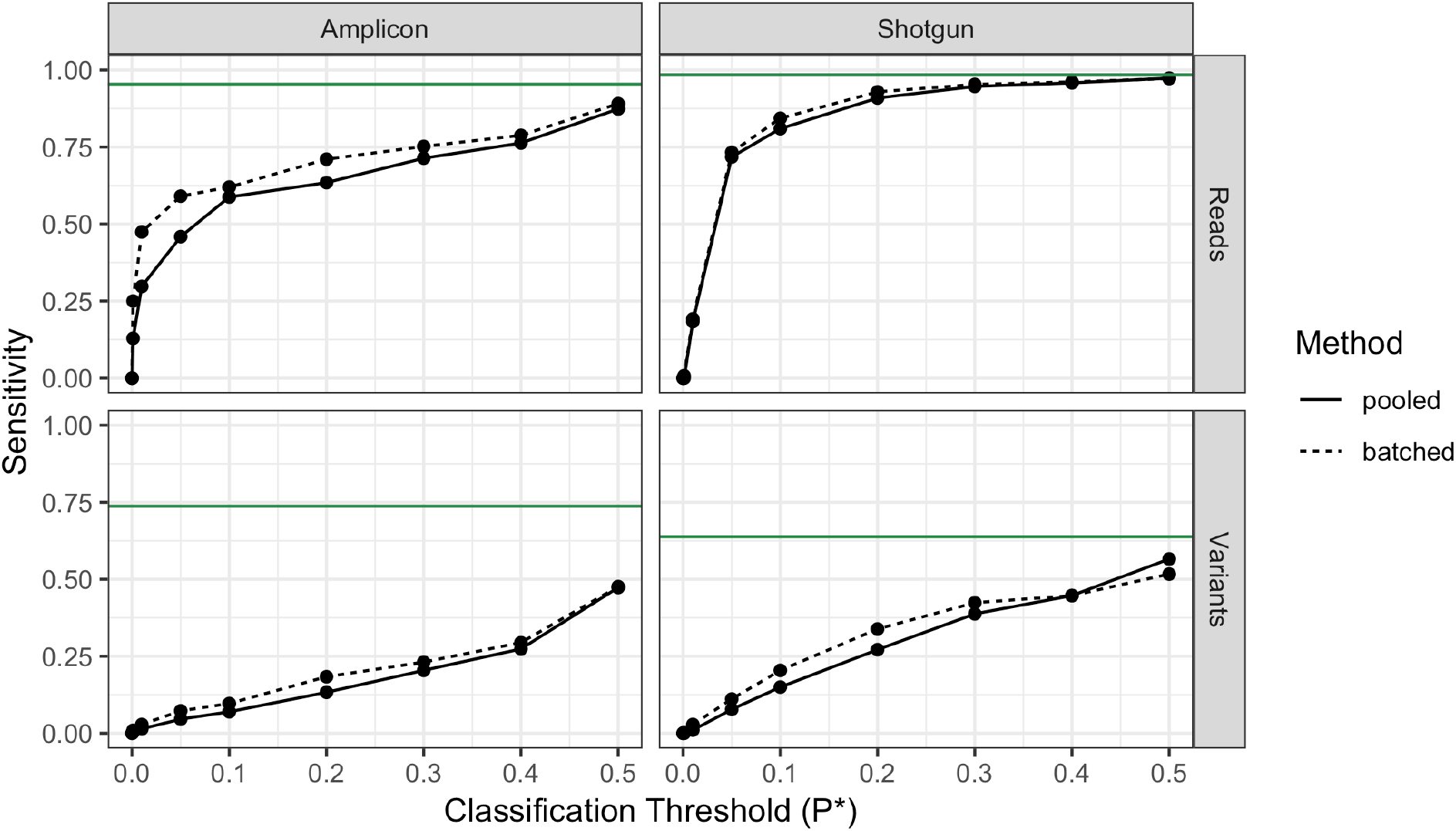
Proportion of contaminants in the *S. bongori* dilution series identified by the decontam frequency method. The frequency method was applied to all data pooled together (solid line), and on a per-batch basis (dashed line). Batches were specified as the sequencing centers for the 16S data, and the DNA extraction kits for the shotgun data. The fraction of contaminants identified (sensitivity) was evaluated on a per-read basis and on a per-variant basis (ASVs for 16S data, genera for shotgun data) over a range of classification thresholds. Green lines show maximum possible classifier sensitivity, given that `decontam` cannot identify contaminants only present in a single sample.

No *Salmonella bongori* reads were classified as contaminants under any P* threshold, and the *S. bongori* variants were assigned the highest scores (>0.98) in these datasets. Negative controls were not included in the shotgun metagenomics sequencing, and only a single negative control was included in the 16S rRNA gene sequencing, so we did not evaluate the prevalence method on this dataset.

Removal of contaminants identified by `decontam` significantly reduced batch effects between sequencing centers and DNA extraction kits (Fig. 5). As the classification threshold increased from P*=0.0 (no contaminants removed) to P*=0.1 (default) to P*=0.5 (aggressive removal), the multi-dimensional scaling ordination distance between samples from different batches decreased. This effect was most dramatic at the intermediate dilutions where both *S. bongori* and various contaminants comprised a significant fraction of the total sequences.

**Figure 5.**
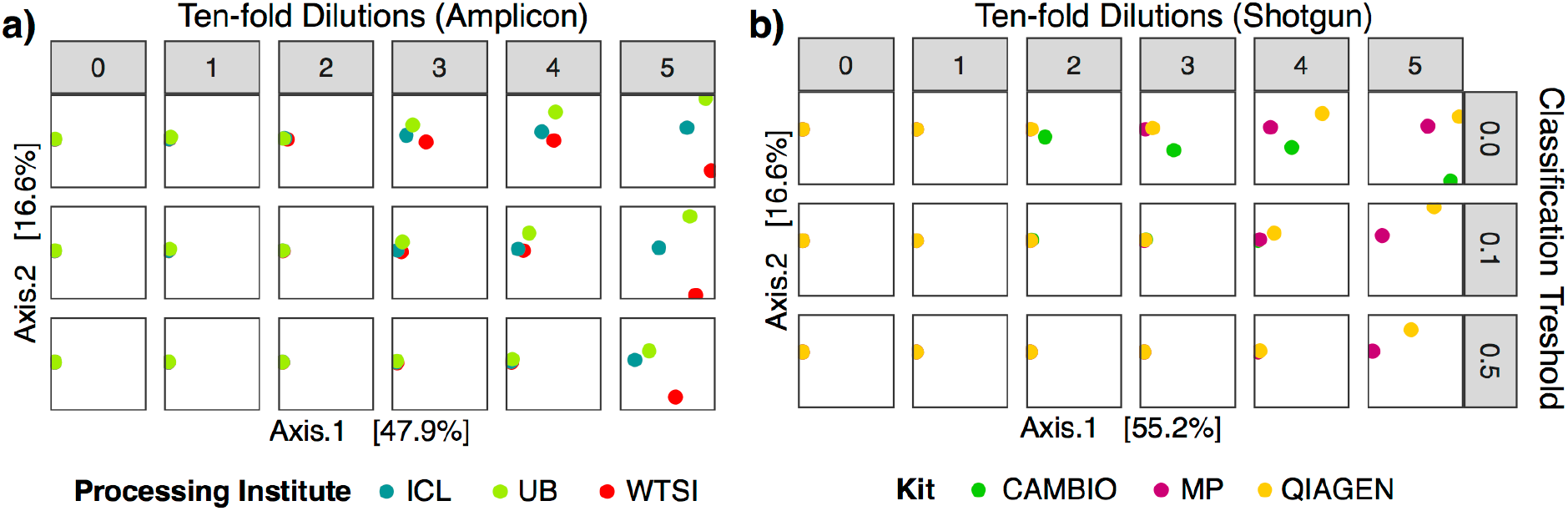
Multi-dimensional-scaling (MDS) ordination of sequenced samples of a monoculture of *S. bongori*, as a function of dilution and the contaminant classification threshold. A dilution series of 0, 1, 2, 3, 4, and 5 tenfold dilutions of a pure culture of *Salmonella bongori* was subjected to **(a)** amplicon sequencing of the 16S rRNA gene at three sequencing centers, and **(b)** shotgun sequencing using three different DNA extraction kits. Contaminants were identified by the `decontam` frequency method with a classification threshold of P*=0.0 (no contaminants identified), P*=0.1 (default), and P*=0.5 (aggressive identification). After contaminant removal, pairwise between-sample Bray-Curtis dissimilarities were calculated, and an MDS ordination was performed. The two dimensions explaining the greatest variation in the data are shown.

In a recent study, Karstens et al. performed an independent evaluation of the `decontam` frequency method on a more complex dilution series constructed from a mock community of 8 bacterial strains. They report that `decontam` correctly classified 74-91% of contaminant reads, and made no false-positive contaminant identifications [57].

### Identification of non-contaminant sequences in a low-biomass environment

Recently, evidence from marker-gene sequencing of placental samples was used to propose that the human placenta harbors an indigenous microbiota [17]. However, contamination has since been proposed as an alternative explanation of those results [15, 44]. To examine this question further, Lauder *et al*. performed 16S rRNA gene sequencing on placenta biopsy samples and multiple negative control samples using two different DNA extraction kits. They found that samples clustered by kit rather than by placental or negative control origin, suggesting that most sequences observed in the placenta samples derived from reagent contamination.

We used the prevalence method as implemented in the *isNotContaminant* function to further explore the possibility that some ASVs in the Lauder *et al*. dataset could be consistent with placental origin, despite being too rare to drive whole-community ordination results. The prevalence score statistic, unlike the frequency score statistic, can be interpreted as a p-value, which allowed us to select candidate non-contaminant ASVs based on a false discovery rate (FDR) threshold [56]. We found that six of the 810 ASVs present in at least five samples were identified as non-contaminants at an FDR threshold of 0.5 (Table 1). Five of those six ASVs matched *Homo sapiens* rather than any bacterial taxa. That is, five of the six ASVs classified by the prevalence method as non-contaminants were classified correctly, as those ASVs were truly present in the placental samples. However, these non-contaminants are not evidence of a placental microbiome, and instead the five *Homo sapiens* ASVs likely arose from off-target amplification of human DNA in the placenta biopsy. The other putative non-contaminant ASV was a Ruminococcaceae variant, a known member of human gut microbial communities [2], but we are unable to establish its ground truth.

**Table 1.**
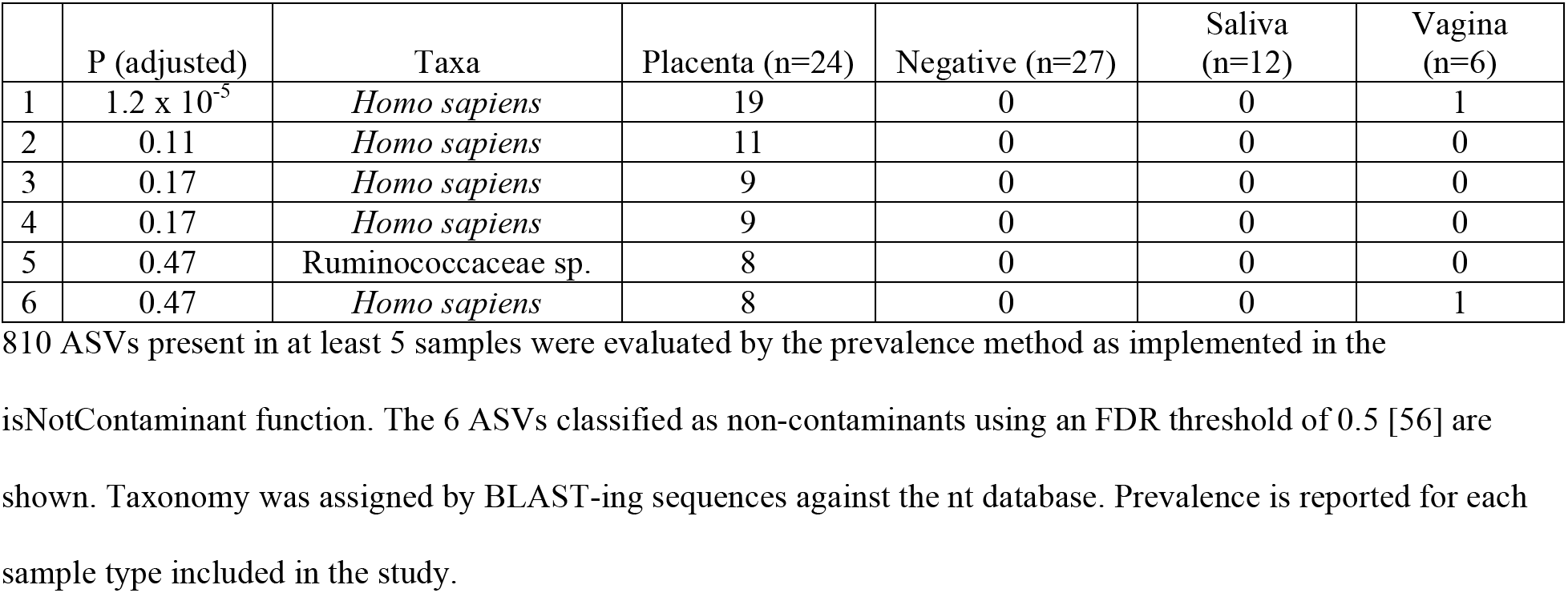
Amplicon sequence variants from placenta samples classified by **decontam** as non-contaminants.

|  | P (adjusted) | Taxa | Placenta (n=24) | Negative (n=27) | Saliva (n=12) | Vagina (n=6) |
| --- | --- | --- | --- | --- | --- | --- |
| 1 | $1.2 \times 10^{-5}$ | *Homo sapiens* | 19 | 0 | 0 | 1 |
| 2 | 0.11 | *Homo sapiens* | 11 | 0 | 0 | 0 |
| 3 | 0.17 | *Homo sapiens* | 9 | 0 | 0 | 0 |
| 4 | 0.17 | *Homo sapiens* | 9 | 0 | 0 | 0 |
| 5 | 0.47 | Ruminococcaceae sp. | 8 | 0 | 0 | 0 |
| 6 | 0.47 | *Homo sapiens* | 8 | 0 | 0 | 1 |

810 ASVs present in at least 5 samples were evaluated by the prevalence method as implemented in the
isNotContaminant function. The 6 ASVs classified as non-contaminants using an FDR threshold of 0.5 [56] are shown. Taxonomy was assigned by BLAST-ing sequences against the nt database. Prevalence is reported for each sample type included in the study.

### Reduction of false-positive associations between the gestational microbiota and preterm birth

A recent exploratory analysis of associations between the gestational vaginal microbiota and preterm birth (PTB) identified a number microbial taxa seemingly associated with PTB [19]. However, the authors concluded that many of these significant associations were run-specific contaminants rather than true biological signal. We used `decontam` to further explore the possibility that some contaminant ASVs were significantly associated with PTB in this dataset.

We generated `decontam` scores using the prevalence, frequency, and combined methods, while specifying the sequencing runs as batches. The scores assigned by all methods showed the expected bimodal score distribution, and the combined method produced the clearest low-score peak (i.e. putative contaminant) (Figure S4). Scores assigned by the frequency and prevalence methods were broadly consistent, especially at low values (Figure S5). We generated a plot similar to the exploratory analysis presented in the original Callahan *et al*. paper, but colored ASVs by the scores assigned by the combined method (Fig. 6). Four of the ASVs most significantly associated with PTB were assigned scores less than 10^−6^, strongly supporting a contaminant origin. Representatives from the genera of those four ASVs have been previously observed as contaminants (*Herbaspirillum:* refs. 12,14,45; *Pseudomonas:* refs. 8,9,12,14,31; *Tumebacillus:* refs. 12,46; *Yersinia:* ref. 47), corroborating decontam’s classification.

**Figure 6.**
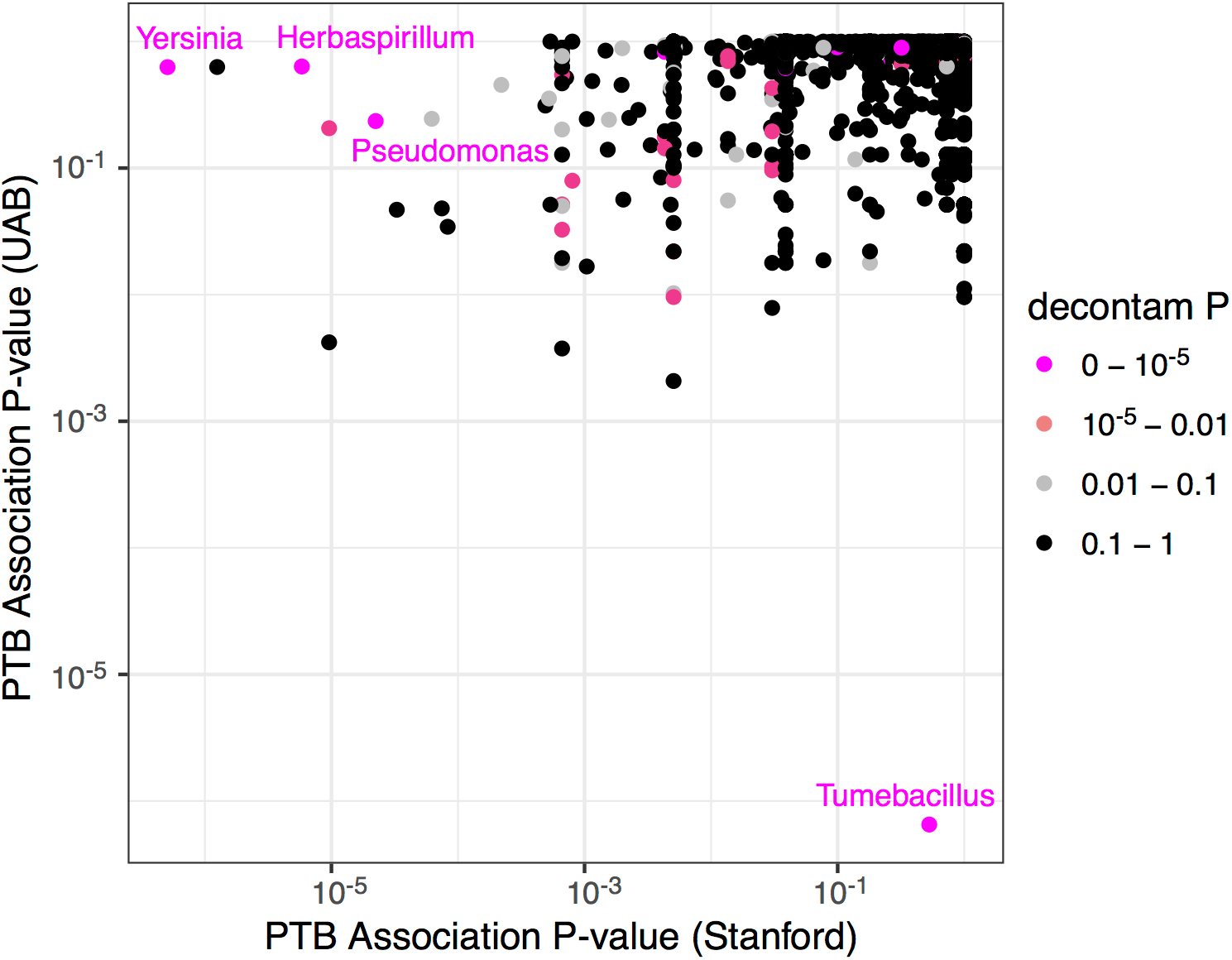
Diagnosing contamination in an exploratory analysis of the vaginal microbiota and preterm birth (PTB). The association between PTB and an increase in the average gestational frequency of various ASVs was evaluated in the two cohorts of women (Stanford and UAB) analyzed in Callahan & DiGiulio *et al*. The x- and y-axes display the P-values of the association between increased gestational frequency and PTB (one-sided Wilcoxon rank-sum test) in the Stanford and UAB cohorts, respectively. Points are colored by the score assigned to them by the *isContaminant* function in the `decontam` R package, using the combined method. Several ASVs that were strongly associated with PTB in either the Stanford or UAB cohorts are clearly identified as contaminants with a `decontam` P value of less than 10^−5^ (genera in magenta text).

## Discussion

Previous work has established two common signatures of contaminants in MGS data: frequency inversely proportional to sample DNA concentration [8,14,16,30], and presence in negative controls [10,31,32]. Building on that work, we developed a simple model of the mixture between contaminant and sample DNA that serves as the basis of frequency-based and prevalence-based statistical classification procedures for identifying contaminants. These methods are implemented in the open source R package decontam, and can be used to diagnose, identify and remove contaminants in marker-gene and metagenomic sequencing datasets.

The classification of contaminants by `decontam` was internally and externally consistent in real datasets. The independent frequency and prevalence methods produced largely consistent results, and the distribution of scores recapitulated the bimodal distribution expected from the proposed mixture model of total DNA. `decontam` classifications were consistent with literature expectations in a subset of genera that the literature unambiguously described as true inhabitants or contaminants.

The classification accuracy of `decontam` increased with the number of samples in which a sequence feature appeared (its prevalence). The rate of false-positive contaminant identification was low for all features, consistent with the findings of an independent benchmarking study [57]. The sensitivity of contaminant identification increased substantially with feature prevalence. As a result, while the specificity and sensitivity of `decontam` on a per-feature basis were sometimes moderate, accuracy evaluated on a per-read basis reached exceptional levels.

In several example datasets, the application of `decontam` improved biological interpretation. Removal of contaminants identified by `decontam` reduced variation due to sequencing center and DNA extraction kit, an oft-cited issue in high-throughput marker-gene and metagenomics studies [14]. `decontam` corroborated recent conclusions that little evidence of an indigenous placenta microbiome existed in a marker-gene dataset from placenta biopsies, and extended that conclusion to rare sequences [15]. `decontam` identified several contaminant taxa in a recent study that a naïve exploratory analysis would have found to be significantly associated with preterm birth [19].

`decontam` improves on current *in silico* approaches to contaminant identification and removal. `decontam` identifies contaminants on a per-sequence-feature basis. `decontam` requires no external knowledge of the pool of potential contaminants. decontam’s statistical classification approach avoids shortcomings of common *ad hoc* threshold approaches. For example, removal of all sequences detected in negative controls also removes abundant true sequences due to cross-contamination among samples [49, 50]. Removal of sequences below an *ad hoc* abundance threshold sacrifices low-frequency true sequences and fails to remove the abundant contaminants most likely to interfere with downstream analysis. In contrast, `decontam` readily detects abundant and prevalent contaminants, while strongly limiting false positives. `decontam` can improve the quality of MGS data and subsequent analyses, at little or no cost to the investigator.

### Using decontam

#### Experimental design

`decontam` is applicable to any MGS dataset for which DNA quantitation data or sequenced negative controls are available. Simple experimental design choices can further improve the performance of decontam. Because reagents contribute significantly to contamination [8, 14–16], negative controls should contain reagents and/or sterile sample collection instruments and should be processed and sequenced alongside true samples. The sensitivity of prevalence-based classification is limited by the number of negative controls, and there is often variation among the contaminants present in each, so we recommend sequencing multiple negative controls. A simulation analysis suggests that 5-6 negative control samples is sufficient to identify most contaminants (assuming a significantly larger number of true samples and a prevalence patterns similar to those seen in the oral dataset, Figure S6), although sensitivity continues to increase with more negative controls. We recommend investigators sequence negative controls for both amplicon and shotgun sequencing approaches, and even if quality checks indicate little or no DNA is present.

In studies large enough to span multiple processing batches (e.g. sequencing runs), we recommend blocking or randomizing samples across the processing batches if possible. Contaminants are often batch-specific, and sample randomization will prevent the conflation of batch-specific contaminants with study outcomes if subsequent contaminant amelioration is not completely effective.

#### Method choice

`The` *isContaminant* function in `decontam` implements distinct frequency- and prevalence-based methods for contaminant identification and can also use both methods in combination. Choice of method should be guided first by the auxiliary data available: frequency-based identification requires DNA quantitation data, and prevalence-based identification requires sequenced negative controls.

DNA concentrations measured from prepared amplicon or shotgun libraries prior to sequencing, often in the form of standardized fluorescent intensities, works effectively with the frequency method in our experience. More effort-intensive methods, such as qPCR, may improve accuracy further if those methods more accurately quantify total DNA [31]. Typically, sufficient variation in DNA concentration for the frequency method to discriminate between contaminants and non-contaminants arises naturally during sample preparation and processing. A positive control dilution series that covers a broad range of input DNA concentrations can guarantee a broad range of sample DNA concentrations [14].

The sensitivity of prevalence-based contaminant identification is limited if few negative controls are sequenced. In very low biomass environments, where contaminant DNA may constitute a majority of sequencing reads, the implementation of the prevalence method in the *isNotContaminant* function can conveniently identify minority non-contaminants.

The score distributions generated by the combined method, which combines the frequency and prevalence scores into a composite score, showed a (slightly) cleaner bimodal distribution than the frequency or prevalence methods alone in the datasets we examinere here (Figs. 2 and S4). Thus, we recommend generating and sequencing negative controls and using the combined approach when the necessary auxiliary data are available, although the frequency and prevalence methods are both independently effective as well.

#### Choice of classification threshold

`decontam` classifies sequence features as contaminants by comparing the score statistic P to a classification threshold P*. We recommend that investigators inspect the distribution of scores assigned by decontam, especially when `decontam` is being applied to large studies spanning multiple batches such as sequencing runs, and consider non-default classification thresholds if so indicated. Typically, an appropriate classification threshold can be read directly off the score histogram. For example, the histogram of scores in Figure 2b showed clear bimodality between very low and high scores, indicating that thresholds in the range from 0.1 to 0.5 would effectively identify the contaminants that make up the low-score mode. In the preterm birth dataset, the low-score mode in the score histogram was much narrower, indicating a threshold of 0.01 would be more appropriate (Fig. S4). Another useful visualization is a quantile-quantile plot of scores versus the uniform distribution.

The scores generated by `decontam` can also be used as quantitative diagnostics instead of as input to a binary classifier. As suggested by our re-analysis of the preterm birth dataset, the `decontam` scores associated with taxa found to be of interest in other analyses can inform subsequent interpretation of the results, and potentially indicate the need for additional confirmatory analyses.

#### Application to heterogeneous samples

`decontam` uses patterns across samples to identify contaminants, but that approach can be less effective when groups of samples have systematically different contaminant patterns. One such scenario arises from separate processing batches that result in batch-specific contaminants. `decontam` allows the user to specify such batches in the data, in which case scores are generated independently within each batch, with the smallest score across batches used for classification by default. Batched classification should be considered when major variation exists in sample processing steps – e.g. different sequencing runs or DNA extraction kits.

The assumptions of decontam, especially the frequency method, can be violated if bacterial biomass systematically differs between groups of samples. For example, if `decontam` were applied to a mixed set of stool (high biomass) and airway (low biomass) samples, real sequences in the airway samples could be classified as contaminants, because they have higher frequency in the low-concentration samples. Therefore, we recommend applying `decontam` independently to samples collected from different environments. Covariation between experimental conditions-of-interest and bacterial biomass could also impinge on the accuracy of contaminant classification, especially if using the frequency method. We have included the *plot condition* function in the `decontam` R package as a convenient way to investigate possible relationships between important experimental conditions and total DNA concentration.

#### Choice of Sequence Feature

`decontam` can be applied to a variety of sequence features derived from MGS data (e.g. OTUs, ASVs, taxonomic groups, MAGs). `decontam` should work most effectively on sequence features that are sufficiently resolved such that contaminants are not grouped with real strains, while also not being overly affected by MGS sequencing noise.

In marker-gene data, we expect the best performance will be achieved with post-denoising ASVs [53, 58] as ASVs are less prone than OTUs to grouping contaminants with related real strains. A general recommendation for metagenomics studies is to use finer taxonomic groups (e.g. species rather than families) and narrower functional categories (e.g. genes rather than pathways).

### Limitations of decontam and Complementary Approaches

`decontam` assumes that contaminants and true community members are distinct from one another. This basic assumption is violated by cross-contamination — contaminant sequences arising from other processed samples [33,49,50]. `decontam` is not designed to remove cross-contamination. MGS studies would benefit from the development of methods to address cross-contamination, and some exciting progress in that area is beginning to be made [33].

`decontam` depends on patterns across samples to identify contaminants, and therefore has low sensitivity for detecting contaminants that are found in very few samples. Since very low-prevalence sequences are often uninformative in downstream analyses, it might often be appropriate to combine `decontam` with the removal of low-prevalence sequences that may be enriched in contaminants that `decontam` did not detect.

### Conclusions

Contaminant removal is a critical but often overlooked step in marker-gene and metagenomics (MGS) quality control [11,14,15,36,46]. Salter *et al*. and Kim & Hofstaedter *et al*. provide excellent pre-sequencing recommendations that reduce the impact of contamination, that can be complemented by *in silico* contaminant removal. Here we introduce a simple, flexible, open-source R package – `decontam` – that uses widely reproduced signatures of contaminant DNA to identify contaminants in MGS datasets. `decontam` requires only data that are in most cases already generated, readily fits into existing MGS analysis workflows, and can be applied to many types of MGS data. Together, our results suggest that `decontam` can improve the accuracy of biological inferences across a wide variety of MGS studies at little or no additional cost.

## Methods

### Oral & control sample processing

Sample DNA was extracted with the PowerSoil®-HTP 96 well Soil DNA Isolation Kit (MO BIO Laboratories, Carlsbad, CA, USA) and then PCR amplified in 2-4 replicate 75-μL reactions using Golay barcoded primers targeting the V4 region of the bacterial 16S rRNA gene [51]. Amplicons were purified, DNA-quantitated using the PicoGreen fluorescence-based Quant-iT dsDNA Assay Kit (ThermoFisher catalog no. Q33120), and pooled in equimolar amounts. After ethanol precipitation and size-selection, amplicons were sequenced in duplicate on two lanes of an Illumina MiSeq v3 flowcell at the W.M. Keck Center for Comparative Functional Genomics (University of Illinois, Urbana-Champaign, USA). Negative controls were processed in parallel with samples beginning at the DNA extraction step (see also Supplemental Methods in Additional File 1).

### Amplicon Sequence Analysis

Duplicate sequencing runs from the oral dataset were concatenated and demultiplexed using QIIME’s split_libraries_fastq.py script [52]. Demultiplexed fastq files from the oral, Salter and placenta datasets were then processed into amplicon sequence variants (ASVs) by DADA2 version 1.8.0 [53]. The final table in the oral dataset consisted of 18,285,750 sequencing reads in 2,420 unique ASVs across 767 samples. Taxonomy was assigned to each sequence using the *assignTaxonomy* function in the dada2 R package [59], and the non-redundant SILVA taxonomic training set (‘silva_nr_v128_train_set.fa’, https://www.arb-silva.de). Further analysis was performed using the phyloseq R package [54].

To improve taxonomic classification accuracy for oral genera, we classified oral sequences a second time using the Human Oral Microbiome Database (HOMD, ref 42, http://www.homd.org/, ‘HOMD_16S_rRNA_RefSeq_V14.5.fasta’). We compared SILVA and HOMD classifications at the genus level, and we resolved assignments for the 80 sequences on which SILVA and HOMD disagreed by NCBI BLAST results.

### Construction of oral and contamination databases

The oral database contains bacterial genera confirmed to inhabit the human oral plaque microbiota by microscopic visualization [55, 39, 41], and genera cultivated from the human oral cavity [42]. These genera are listed with their literature citations in the ‘oral_database.csv’ file and the HOMD [42] as ‘named’ or ‘unnamed’ entries. The contamination database contains bacterial genera previously reported as contaminants in 16S rRNA gene negative controls. Contaminant genera are listed with their literature citations in the ‘contamination_database.csv’ file. Genera were categorized into three groups by comparison to the oral and contamination databases: Contaminant, if present in the contamination database and not in the oral database; Oral, if present in the oral database and not in the contamination database, and Ambiguous otherwise. Reference-based classification of ASVs was performed based on their assigned genus, and if no genus was assigned then the ASV was classified as Ambiguous.

## List of abbreviations

MGS: marker-gene and metagenomic sequencing
ASV: amplicon sequence variant
MAG: metagenome-assembled genome
OTU: operational taxonomic unit

## Declarations

### Ethics approval and consent to participate

The study was approved by the Administrative Panels on Human Subjects Research (Stanford IRB protocol #21586), and by the Human Research Protection Program at UCSF (UCSF IRB protocol ##11-06283). All research subjects provided written informed consent prior to specimen collection.

### Consent for publication

Not applicable.

### Availability of data and material

The sequencing data from the human oral microbiome dataset are available at the SRA under SRA accession number SRP126884. The R markdown analysis scripts and necessary input data are available at https://benjjneb.github.io/DecontamManuscript/

## Supplementary Information

Supplemental methods, figures, and tables can be found in Additional File 1.

## Authors’ contributions

D.A.R. acquired funding; N.M.D., D.M.P., S.P.H., D.A.R., and B.J.C. designed research; N.M.D., D.M.P., and B.J.C. performed research; N.M.D., D.M.P., and B.J.C. analyzed data; N.M.D. and B.J.C. wrote the manuscript; N.M.D., D.M.P., S.P.H., D.A.R., and B.J.C. critically revised the manuscript. All authors approved the final version.

## Acknowledgments

We thank the study participants for specimen donation, Dr. Ava Wu and Danielle Drury at the UCSF School of Dentistry for collecting oral mucosa specimens, Dr. Mark Ryder at UCSF for managing the UCSF protocol, and Dr. Les Dethlefsen and members of the Relman Lab for helpful discussions.

